# Targeted Cleavage and Polyadenylation of RNA by CRISPR-Cas13

**DOI:** 10.1101/531111

**Authors:** Kelly M. Anderson, Pornthida Poosala, Sean R. Lindley, Douglas M. Anderson

## Abstract

Post-transcriptional cleavage and polyadenylation of messenger and long noncoding RNAs is coordinated by a supercomplex of ~20 individual proteins within the eukaryotic nucleus^1,2^. Polyadenylation plays an essential role in controlling RNA transcript stability, nuclear export, and translation efficiency^3–6^. More than half of all human RNA transcripts contain multiple polyadenylation signal sequences that can undergo alternative cleavage and polyadenylation during development and cellular differentiation^7,8^. Alternative cleavage and polyadenylation is an important mechanism for the control of gene expression and defects in 3’ end processing can give rise to myriad human diseases^9,10^. Here we show that fusion of catalytically dead Cas13 to a single mammalian polyadenylation factor, Nudix hydrolase 21 (NUDT21), allows for site-specific CRISPR-Cas13-guided cleavage and polyadenylation of RNA in mammalian cells. This approach, which we named *Postscriptr*, can be utilized for the non-genomic manipulation of gene expression and may have potential future therapeutic applications for treating human RNA processing diseases.

---

The 3’ site of RNA cleavage and addition of a poly(A) tail is precisely determined by intrinsic polyadenylation signal (PAS) sequences typically composed of a canonical hexamer motif AAUAAA and upstream and downstream sequence elements (USE and DSE, respectively)^11^. The AAUAAA PAS motif is found ~25 nucleotides (nts) upstream of the RNA cleavage site and is directly bound by two components of the cleavage and polyadenylation specific factor (CPSF) complex: cleavage and polyadenylation factor 30 (CPSF30) and WD repeat-containing protein 33 (WDR33)^12^. Components of the cleavage factor Im (CFIm) complex bind directly to the USE motif UGUA which occurs ~50 nts upstream of the RNA cleavage site^8^. In transcripts that lack the canonical AAUAAA motif, the CFIm complex functions as the primary determinant of poly(A) signal recognition^13^. Nudix Hydrolase 21 (NUDT21/CPSF5), the RNA binding component of the CFIm complex, functions as an activator of 3’ end processing and regulator of alternative polyadenylation site choice^14^. Binding of the CFIm complex is among the first steps in 3’ end processing and functions to recruit additional co-factors to the 3’ end processing machinery^15,16^. Direct interactions between components of the polyadenylation supercomplex and the C-terminal domain of RNA polymerase II trigger the disassembly of the elongation complex and coordinate 3’ end processing with transcription termination^17^.

Bacterial-derived Type VI CRISPR-Cas systems encode a family of RNA-guided endoribonucleases, named Cas13^18–20^. Similar to Cas9, mutation of residues within the nuclease domains of Cas13 generate a catalytically dead enzyme that retains RNA binding affinity (dCas13). Recently, fusion of dCas13 to mammalian RNA modifying enzymes has been shown to be useful for manipulating RNA in mammalian cells^21–23^. To direct polyadenylation complex formation using dCas13, we designed three fusion proteins combining the catalytically dead Type VI-B Cas13 enzyme from *Prevotella sp. P5–125* (dPspCas13b) with RNA binding components of the human 3’ end processing machinery, including CPSF30, WDR33, and NUDT21 (Fig. 1a). We included an N-terminal 3x FLAG epitope tag for protein detection and a long flexible peptide linker [GGGGSGGGGS] between dPspCas13b and the polyadenylation components to reduce the likelihood of steric hindrance.

**Figure 1.**
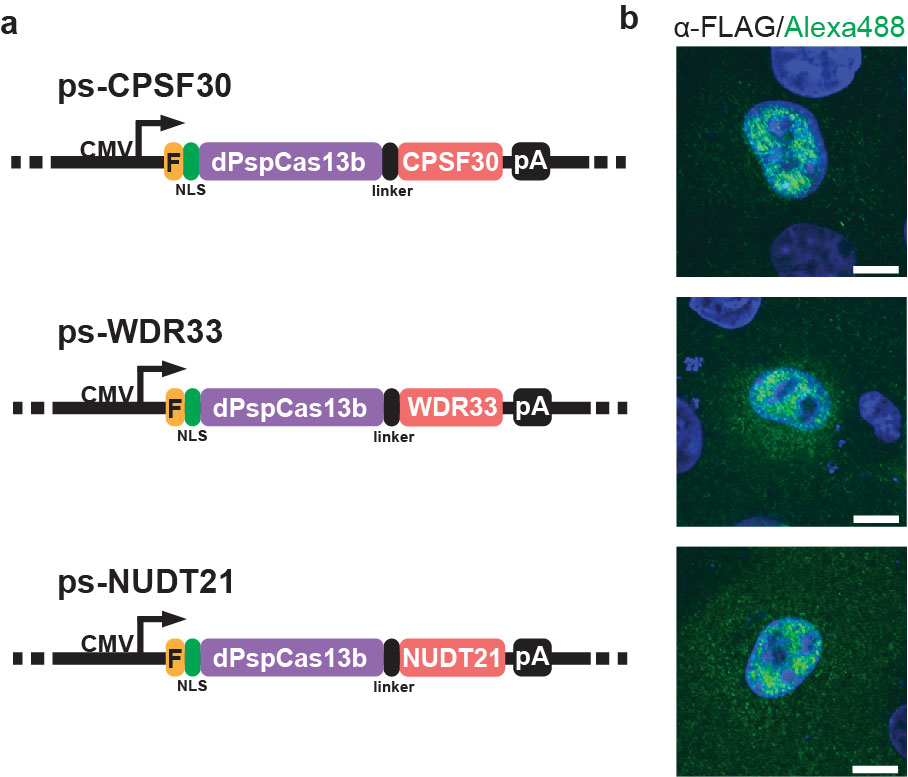
Design and expression of fusion proteins encoding catalytically dead Cas13 and human polyadenylation complex factors. (**a**) Diagram of the vectors encoding fusions between catalytically dead PspCas13b and components of the mammalian polyadenylation supercomplex, CPSF30, WDR33, and NUDT21. F – 3xFLAG epitope; NLS – Ty1 nuclear localization sequence; pA – SV40 polyadenylation sequence. (**b**) Immunohistochemistry using a primary anti-FLAG antibody and an Alexa488 conjugated secondary antibody detecting the nuclear localization of the dPspCas13b fusion proteins expressed in mammalian COS7 cells. Scale bars, 10 μm. Results are representative of 3 independently performed experiments.

While previous dCas13-mediated RNA editing applications occurred within the cytosol, post-transcriptional cleavage and polyadenylation of RNA occurs within the nucleus and therefore requires efficient nuclear localization of dCas13 fusion proteins. The large size of dCas13 proteins and their lack of intrinsic nuclear localization signals (NLSs) could prevent their efficient nuclear localization in mammalian cells. Consistent with this, dPspCas13b fusion proteins lacking a mammalian NLS were retained in the cytoplasm when expressed in mammalian COS7 cells, as detected by immunocytochemistry using an anti-FLAG antibody (Supplementary Fig. 1). Surprisingly, the addition of a classical SV40 NLS or bipartite NLS from nucleoplasmin (NPM) did not promote efficient nuclear localization of dPspCas13b fusion proteins (Supplementary Fig. 1). However, the addition of a single copy of the non-classical bipartite NLS derived from the yeast Ty1 retrotransposon, which utilizes normal cellular protein import machinery but contains a linker sequence nearly three times as long as a typical bipartite NLS^24,25^, resulted in robust nuclear localization of all three dPspCas13b fusion proteins (Fig. 1b).

We designed a reporter gene to detect alternative cleavage and polyadenylation (APA) in mammalian cells capable of switching between fluorescent and non-fluorescent open reading frames of superfolder GFP (sfGFP) (Fig. 2). sfGFP forms a beta barrel comprised of 11 antiparallel beta strands, which can tolerate sequence insertions between the 10^th^ and 11^th^ beta strands but loses fluorescence if the 11^th^ beta strand is removed^26–28^. We generated an sfGFP reporter construct with the coding sequence for the 11^th^ beta strand embedded within a prototypical mammalian intron (second intron of the rabbit beta globin gene) (sfGFPapa) (Fig. 2a). The coding sequence of the 11^th^ beta strand was designed in-frame with the upstream sfGFP coding sequence so that translation of the proximal open reading frame would encode a complete sfGFP sequence, albeit with a 14 amino acid linker sequence between the 10^th^ and 11^th^ beta strand resulting from translation of the intervening intronic sequence [sfGFP(1-10)-L-11]. Expression of the sfGFPapa reporter in mammalian COS7 cells resulted in almost no detectable fluorescence, suggesting efficient removal of the intron containing the 11^th^ beta strand and translation of the open reading frame encoding the non-fluorescent sfGFP(1-10) protein (Fig. 2b-d). However, cells expressing the sfGFPapa reporter treated with the splicing inhibitor isoginkgetin^29^ resulted in detectable green fluorescence after 24 hours, demonstrating that the modified sfGFP open reading frame containing the linker between the 10^th^ and 11^th^ beta strands is functional (Supplementary Fig. 2a,b). Insertion of an SV40 polyadenylation cassette downstream of coding sequence for the 11^th^ beta strand of sfGFP (sfGFPapa-pA) resulted in robust green fluorescence, demonstrating that intronic cleavage and polyadenylation of the sfGFPapa reporter promotes utilization of the functional sfGFP(1-10)-L-11 open reading frame (Fig. 2e-h). Together, these data demonstrate that the sfGFPapa reporter gene encodes both functional and non-functional open reading frames which can be switched in response to changes in splicing or intronic cleavage and polyadenylation.

**Figure 2.**
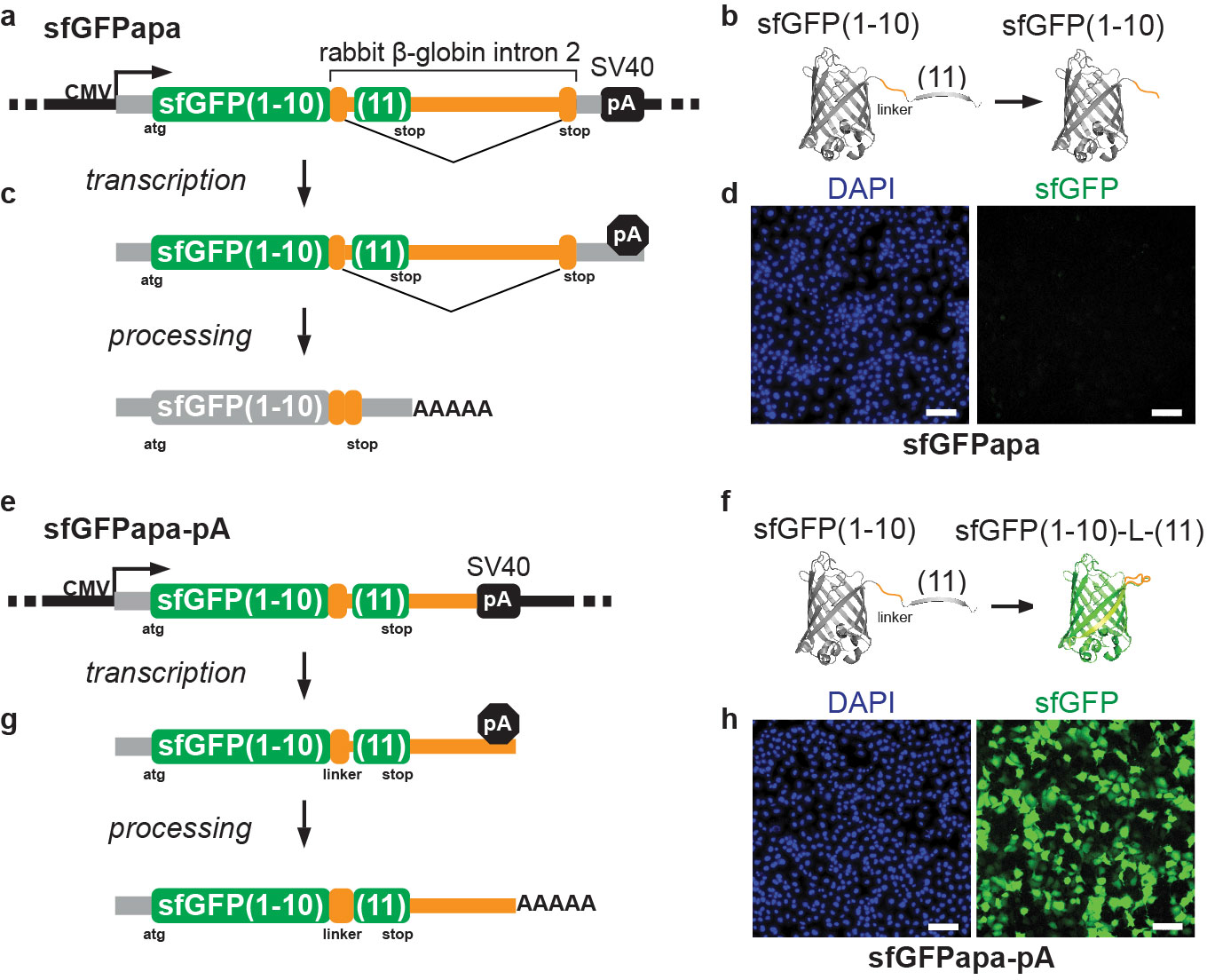
Design and Generation of a Fluorescent Reporter for RNA Cleavage and Polyadenylation in Mammalian Cells. (**a**) Diagram of the sfGFPapa reporter gene. (**b**) The green fluorescent protein superfolder GFP (sfGFP) forms a beta barrel composed of 11 beta strands. Removal of the 11^th^ beta strand abolishes fluorescence. (**c**) Transcription of the sfGFPapa reporter and processing of the resulting transcript removes the coding sequence of the 11^th^ sfGFP beta strand. (**d**) Removal results in a lack of fluorescent signal in mammalian cells. (**e**) Diagram of the sfGFPapa-pA reporter plasmid, which contains an upstream polyadenylation sequence. (**f**) sfGFP can tolerate linker sequences between the 10^th^ and 11^th^ beta strands without abolishing fluorescence. (**g**) Transcription of the sfGFPapa-pA reporter and processing of the resulting transcript. (**h**) Processed transcripts result in robust green fluorescence due to translation of the sfGFP(1-10)-L-(11) functional open reading frame. Scale bars in (**d**) and (**h**), 100 μm. Results are representative of 3 independently performed experiments.

To determine whether the dPspCas13b fusion proteins could promote CRISPR-Cas13-mediated cleavage and polyadenylation of the sfGFPapa reporter mRNA, we designed a crRNA targeting an intronic sequence downstream of the coding sequence of the 11^th^ beta strand of sfGFP in the sfGFPapa reporter (Fig. 3a). In live cell assays, no fluorescence signal was detected in cells expressing any of the dPspCas13b fusion proteins using a non-targeting crRNA, relative to the sfGFPapa reporter alone (Fig. 3b, Supplementary Fig. 3). However, expression of the intron targeting crRNA with the dPspCas13b-NUDT21 fusion protein, but not the CPSF30 or WDR33 fusions proteins, resulted in detectable green fluorescent cells after 24 hours (Fig. 3b, Supplementary Fig. 3).

**Figure 3.**
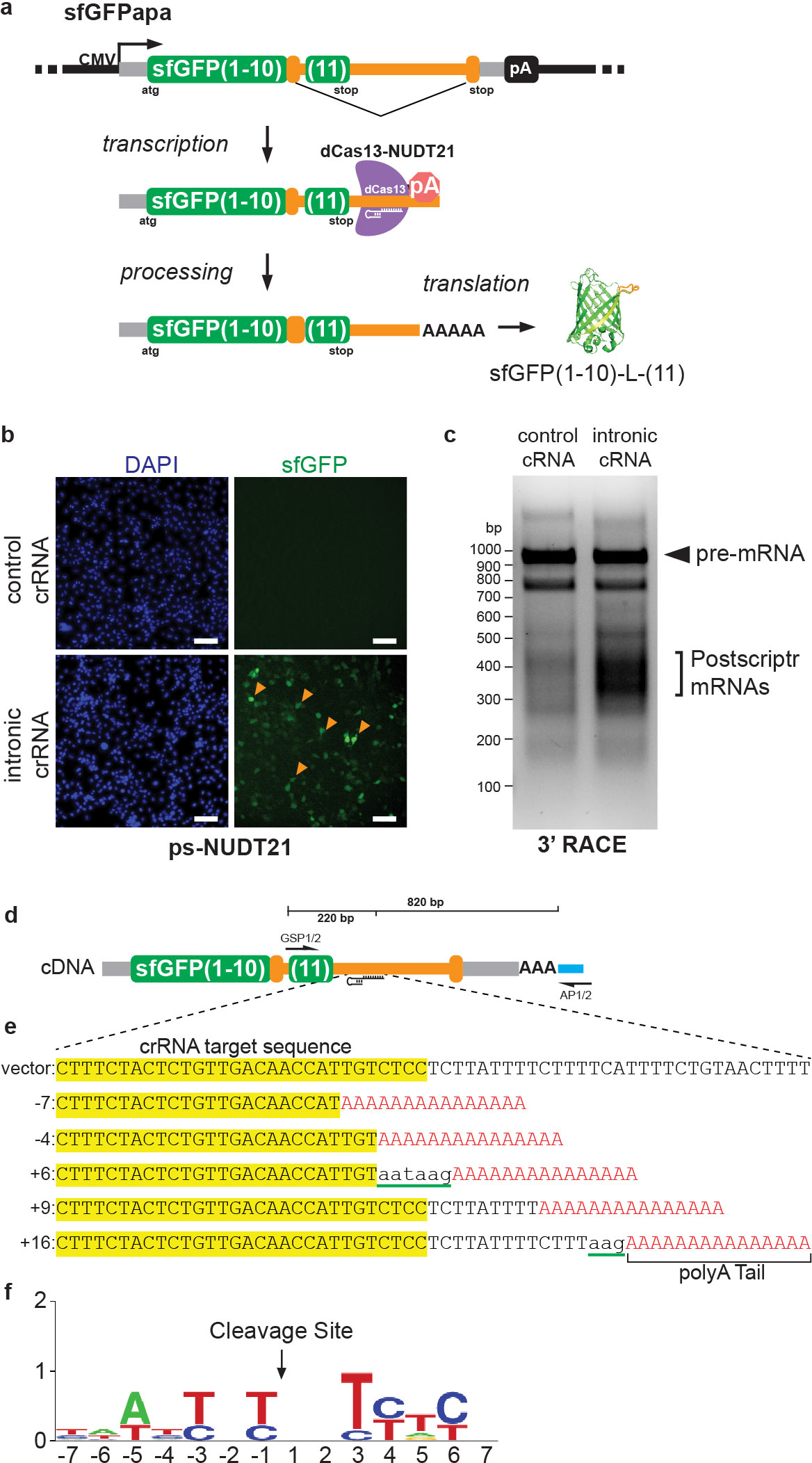
CRISPR-Cas13-mediated Cleavage and Polyadenylation of a Reporter mRNA in Mammalian Cells. (**a**) Diagram showing the *Postscriptr* targeting strategy to induce alternative cleavage and polyadenylation of the sfGFPapa reporter construct. A crRNA was designed to target an intronic sequence downstream of the coding sequence of the 11^th^ beta strand of sfGFP. (**b**) Expression of the dPspCas13b-NUDT21 fusion protein and intronic targeting crRNA resulted in green fluorescent cells after 24 hours. (**c**) 3’RACE products amplified from cells expressing dPspCas13b-NUDT21 fusion protein and non-targeting control or intronic targeting crRNA. (**d,e**) Sequences of the five most proximal 3’RACE products relative to the crRNA target site. Please see Table 1 for complete list of 3’RACE sequences. Green underlined and lowercase nucleotides highlight 3’ non-templated nucleotide addition. (**f**) Nucleotide frequencies at *Postscriptr*-mediated cleavage and polyadenylation sites.

We performed 3’ rapid amplification of cDNA ends (3’ RACE) to determine whether the fluorescence signal was due to CRISPR-Cas13-mediated targeted cleavage and polyadenylation of the sfGFPapa reporter mRNA. RACE compatible cDNA was generated from total RNA using a poly(T) oligonucleotide containing two 5’ nested primer sequences. Using two nested reporter specific primers upstream of the crRNA target sequence, we detected a high molecular weight band corresponding to the predicted size of the sfGFPapa pre-mRNA transcript for both samples (Fig. 3c). Interestingly, in cells targeted with the intronic crRNA, we detected an additional broad band of smaller RACE products, suggesting that cleavage and polyadenylation may have occurred near the targeted crRNA sequence (Fig. 3c,d). To determine the sequence of the 3’ RACE products, we gel purified the 200-500 base pair region and subcloned them using TOPO TA cloning. We isolated and sequenced 11 unique clones which revealed that sfGFPapa mRNAs were cleaved and polyadenylated at sites ranging from −7 to +110 nts relative to the 3’ side of the crRNA target sequence (Fig. 3e). In contrast to mammalian cleavage which occurs primarily after a CA dinucleotide, *Postscriptr*-induced cleavage most often occurred at a C or T nucleotide (Fig. 3f). Interestingly, four of the 11 clones showed a 3’ addition of non-templated nucleotides prior to poly (A) tail elongation, which is thought to be rare in mammalian species but observed in plants^30^. These data demonstrate that the dPspCas13b-NUDT21 fusion protein was sufficient to induce cleavage and polyadenylation of a reporter mRNA at an intronic sequence targeted by a crRNA.

To determine whether *Postscriptr* could promote alternative cleavage and polyadenylation of an endogenously expressed human mRNA, we targeted transcripts encoding the human sterol regulatory element binding protein 1 (SREBP1). SREBP1 is a ubiquitously expressed transcription factor which transactivates genes that contain sterol regulatory elements (SREs) and encode proteins controlling lipid synthesis and uptake, such as the low density lipoprotein receptor (*LDLR*) (Fig. 4)^31^. SREBP1 is first synthesized as an inactive precursor protein anchored to the membrane of the endoplasmic reticulum (ER)^32^ (Fig. 4a). In sterol depleted cells, proteolytic cleavage of SREBP1 by S2P liberates an N-terminal DNA-binding fragment that allows SREBP1 to enter the nucleus and activate gene transcription^33^ (Fig. 4a). We note that *SREBP1* transcripts in the liver can utilize an endogenous intronic PAS and generate a protein fragment terminated at a site adjacent to the transcriptionally active S2P cleavage product [SREBP1aΔ, AB373958; SREBP1cΔ, AB373959] (Fig. 4b,c). We were interested in whether *Postscriptr* could promote the utilization of the SREBP1 intronic PAS in a non-liver cell line and upregulate *SREBP1* target gene expression. We co-expressed the dPspCas13b-NUDT21 fusion protein with either an *SREBP1*-targeting or non-targeting crRNA in human HEK293T cells (Fig. 4d). Performing 3’ RACE, we detected a strong band corresponding to the approximate size of the predicted 3’RACE product in cells transfected with the *SREBP1*-targeting crRNA, and no bands were detected in cells transfected with the control crRNA (Supplementary Fig. 4). Direct sequencing of this 3’ RACE product revealed that *SREBP1* transcripts were cleaved and polyadenylated 71 nts downstream of the crRNA target sequence and that splicing of the terminal exon was inhibited (Fig. 4d,e). Cleavage occurred at a CA dinucleotide 36 nts downstream of the reported cleavage site in humans and generated an open reading frame which terminated at a stop codon within the retained intron sequence, adding an additional 16 amino acids to the C-terminus of SREBP1 (Fig. 4e,f). Gene expression analyses in a mixed population of *SREBP1*-targeted and non-targeted HEK293T cells showed that the overall *SREBP1* transcript levels were unchanged when using qRT-PCR and PCR primers upstream of the crRNA target sequence (Fig. 4g). However, primers spanning the targeted exon showed a significant decrease in *SREBP1* transcript levels in targeted versus non-targeted cells, suggesting that transcription termination may be coupled with *Postscriptr*-mediated cleavage and polyadenylation (Fig. 4h). Interestingly, in *SREBP1*-targeted cells, we further detected a significant increase in transcript levels of the well-characterized SREBP-responsive gene, *LDLR* (Fig. 4i).

**Figure 4.**
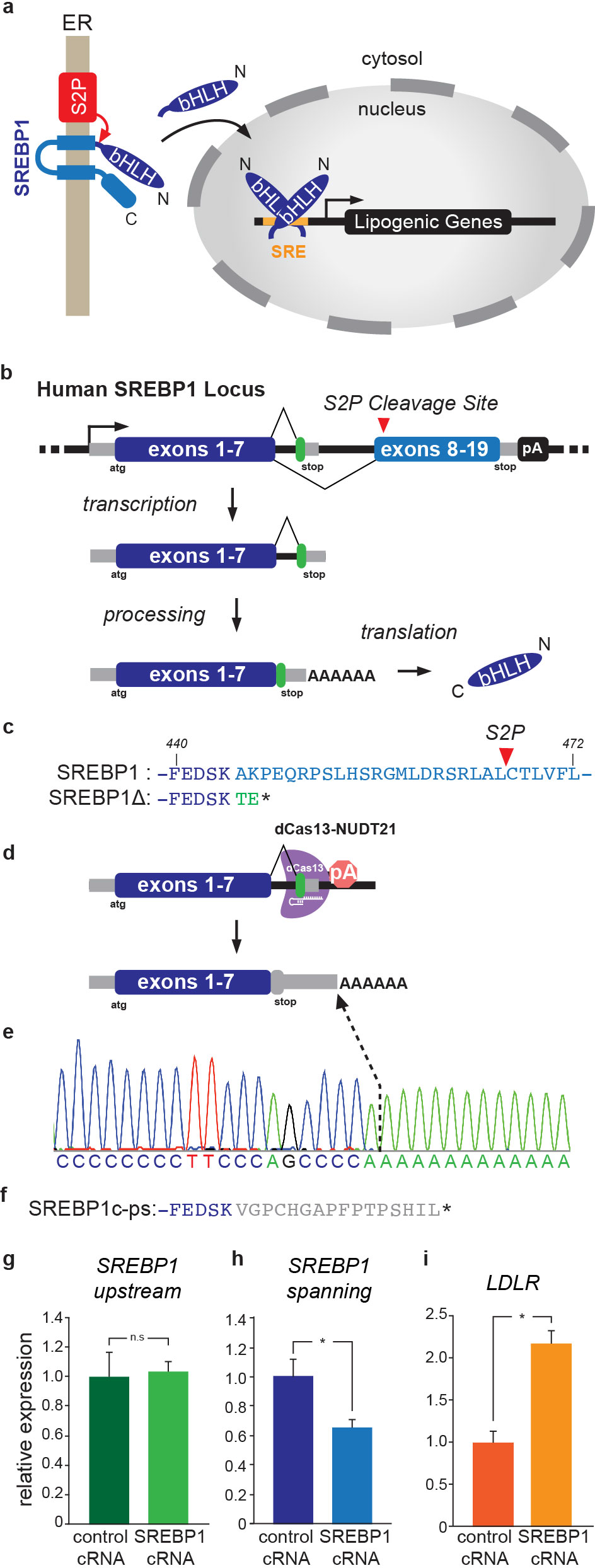
*Postscriptr*-mediated Alternative Cleavage and Polyadenylation of human SREBP1 in HEK293T cells. (**a**) Diagram of the SREBP1 maturation pathway. (**b**) The human SREBP1 locus contains an intronic PAS between exon 7 and exon 8, which results in translation of an SREBP1Δ isoform which terminates translation adjacent to the normal S2P cleavage site (**c**). (**d**) Diagram of the *Postscriptr*-mediated targeting strategy and sequence of the recovered *Postscriptr*-induced cleaved and polyadenylated transcript (**e**) and predicted translational stop (**f**). Quantitative realtime-PCR (qRT-PCR) gene expression analysis of SREBP1 transcript levels upstream of the crRNA target sequence (**g**), spanning the crRNA target sequence (**h**) and transcript levels of the LDLR gene (**i**) in HEK293T cells targeted with a non-targeted or SREBP1-targeted crRNA.

In conclusion, we have developed a unique CRISPR-Cas13 RNA modifying technology to induce site-specific cleavage and polyadenylation of RNA in mammalian cells. This approach utilizes a dPspCas13b-NUDT21 fusion protein, which we have shown requires a non-classical bipartite NLS for nuclear localization in mammalian cells. We demonstrated that the dPspCas13b-NUDT21 fusion protein was sufficient to promote cleavage and polyadenylation at an intronic sequence within a novel gain-of-function fluorescent reporter, sfGFPapa. The ability of *Postscriptr* to promote cleavage and polyadenylation of the sfGFPapa reporter was striking, given that the second intron of the rabbit beta globin gene does not contain any known endogenous PAS sequences. These findings show that NUDT21 can facilitate the recruitment of a functional 3’ end processing complex at a crRNA targeted sequence. While only 25 kDa in size, NUDT21 forms a number of direct protein interactions with key RNA processing components, including Poly(A) Polymerase^34^. These findings support the role of NUDT21 as an emerging master regulator of alternative cleavage and polyadenylation site choice in mammals^35,36^.

We further showed that *Postscriptr* could direct cleavage and polyadenylation at an endogenous intronic PAS of the human *SREBP1* gene. For both the sfGFPapa reporter and SREBP1, the dPspCas13b-NUDT21 fusion protein induced cleavage and polyadenylation 3’ to the crRNA target sequence. However, it is currently unclear why editing of the intronic SREBP1 PAS resulted in a single cleavage site, whereas cleavage of the sfGFPapa reporter occurred across a range of ~100 nts. We speculate that intrinsic PAS flanking sequences may be required to help guide cleavage site specificity. Future studies are necessary to delineate the minimal upstream and downstream sequence elements which may enhance the precision of *Postscriptr* targeting at ectopic target sites.

It is currently unclear why fusion of dPspCas13b to either CPSF30 or WDR33 were not sufficient to promote cleavage and polyadenylation of the sfGFPapa reporter in our assays. Since both CPSF30 and WDR33 have been shown to make direct contacts with the AAUAAA PAS motif, one possibility may be a requirement of the PAS RNA to promote a ternary complex formation among these components. Alternatively, tethering of CPSF30 and WDR33 may alone be insufficient to induce 3’ end processing, a possibility consistent with data showing that insertion of a minimal synthetic PAS (including an AAUAAA motif and downstream sequence elements) within the second intron of a rabbit beta globin gene reporter was insufficient to promote intronic cleavage and polyadenylation^37^.

While the endonuclease activity of Cas13 is highly effective for gene knockdown in mammalian cells^21^, *Postscriptr* technology allows for a unique approach to manipulate RNA which may be useful for the study of 3’ end RNA processing and for the manipulation of endogenous gene expression. Here we have utilized this technology to manipulate the alternative cleavage and polyadenylation at intronic pre-mRNA transcript sequences, however this approach could be applied to other mRNA transcript regions or different RNA species. One interesting example would be the interrogation of lncRNA function, many of which control gene expression through recruitment of chromatin modifying complexes or by establishing permissive local chromatin environments^38–41^. *Postscriptr*-guided cleavage, polyadenylation, and transcriptional termination of lncRNAs could allow for the identification of functional lncRNA domains or functional genomic regions. Lastly, numerous loss-of-function mutations identified within poly(A) signal sequences have been shown to result in abnormal RNA processing and can cause human disease^42–44^. Here, future therapeutic applications of *Postscriptr* technology may be a useful approach for the rescue of normal RNA cleavage and polyadenylation site choice.

## ACKNOWLEDGEMENTS

We thank Dr. Lynne Maquat for comments on the manuscript and Leisha Manchin for technical assistance. pC0043-PspCas13b crRNA backbone was a gift from Feng Zhang (Addgene plasmid # 103854; http://n2t.net/addgene:103854; RRID:Addgene_103854). pC0049-EF1a-dPSPCas13b-NES-HIV, H133A/H1058A was a gift from Feng Zhang (Addgene plasmid # 103865; http://n2t.net/addgene:103865; RRID:Addgene_103865).

## AUTHOR CONTRIBUTIONS

K.M.A. and D.M.A. designed the experiments. K.M.A., P.P., S.R.L and D.M.A performed the experiments and analyzed data. K.M.A. and D.M.A generated the figures and wrote the manuscript.

## AUTHOR INFORMATION

The authors declare no competing financial interests. Correspondence and requests for materials should be addressed to D.M.A.

## METHODS

### Synthetic DNA and Cloning

The coding sequences of human CPSF30, WDR33 and NUDT21were designed and synthesized for assembly as gBlocks (IDT, Integrated DNA Technologies) and cloned as C-terminal fusions to dPspCas13b into a modified version of the CS2 mammalian expression vector containing a 5’ T7 promoter (CSX). The sfGFPapa reporter was designed and synthesized as two separate gBlock gene fragments and subcloned into the CS2 mammalian expression vector. Guide RNAs were designed antisense to pre-mRNA sequences of the sfGFPapa reporter or human *SREPB1* transcripts and were cloned into pC00043^22^. All guide RNAs were designed to be 30 nts in length and start with a 5’ G. All plasmid insert sequences were fully verified by Sanger sequencing. Please see Supplementary Table 1 for fusion protein and reporter sequences.

### Cell Lines, Transient Transfections, and Drug Treatments

The COS7 cell line was maintained in DMEM supplemented with 10% Fetal Bovine Serum (FBS) with penicillin/streptomycin at 37°C in an atmosphere of 5% CO_2_. Cells were seeded and transiently transfected using Fugene6 (Promega) according to manufacturer’s protocol. Twenty-four hours after transfection, cells were either imaged, or treated with DMSO or Isoginkgetin and cultured for an additional 24 hours prior to imaging. For live cell imaging, cells were washed twice with DPBS, counterstained with 10 μg/ml Hoechst 33342 (Invitrogen) in DPBS for 10 minutes at room temperature and imaged on a ZOE Fluorescent Cell Imager (Bio-Rad).

### Antibodies and Immunohistochemistry

Transiently transfected COS7 cells were fixed in 4% formaldehyde in PBS for 20 minutes, blocked in 3% Bovine Serum Albumin (BSA) and incubated with primary anti-FLAG (Sigma, F1864) at 1:1000 in 1% BSA for 1 hour at room temperature. Cells were subsequently incubated with an Alexa488 conjugated secondary antibody (Thermofisher) in 1% BSA for 30 minutes at room temperature. Coverslips were mounted using anti-fade fluorescent mounting medium containing DAPI (Vector Biolabs, H-1200) and imaged using confocal microscopy.

### 3’ Rapid Amplification of cDNA Ends (3’ RACE)

Transfected cells were washed twice with DPBS and harvested in 1 ml of Trizol to isolate total RNA. RACE compatible cDNA was generated using SuperScript III (Invitrogen) according to manufacturer’s protocol, with the exception of the addition of a custom oligo d(T) oligonucleotide containing two 5’ nested primer sequences. RACE PCR was performed using two nested target gene specific primers and touchdown PCR protocol as follows: 94°C for 1 minutes (1 cycle); 94°C for 30 seconds, 72°C for 1 minute (5 cycles); 94°C for 30 seconds, 70°C for 4 minutes (5 cycles); 94°C for 20 seconds, 68°C for 1 minute (25 cycles); 72°C for 5 minutes. Please see Supplementary Table 1 for a list of qRT-PCR primers and 3’RACE PCR primers.

## SUPPLEMENTARY DATA

### SUPPLEMENTARY FIGURES

**Supplementary Figure 1.**
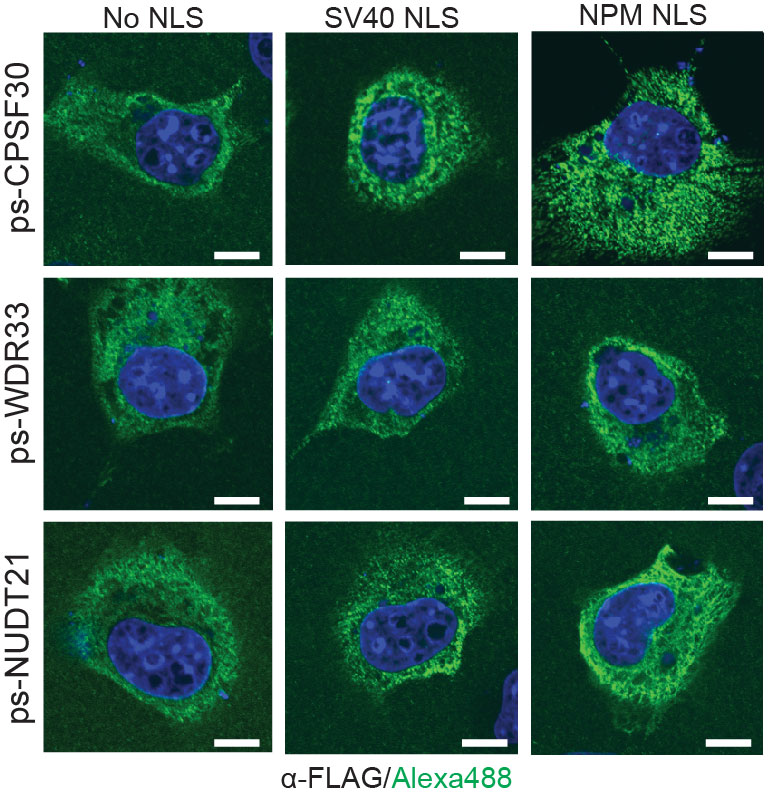
Classic mammalian nuclear localization signals were insufficient to promote nuclear localization of dPspcas13b fusion proteins. Immunohistochemistry using a primary anti-FLAG antibody and secondary Alexa488 conjugated secondary antibody were used to detect the localization of dPspCas13b fusion proteins expressed in mammalian COS7 cells. Fusion proteins contained either no NLS, a classic SV40 NLS or the bipartite NLS from Nucleoplasmin (NPM). Scale bars, 10 μm.

**Supplementary Figure 2.**
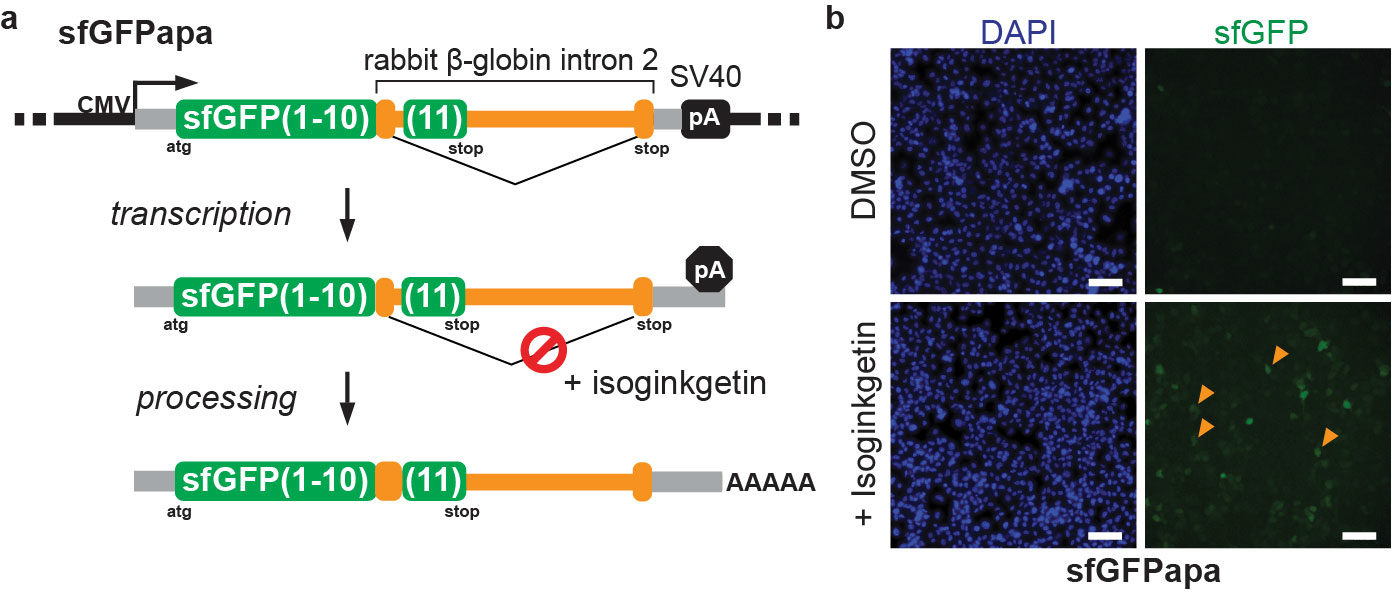
Inhibiting splicing promotes expression of the sfGFP(1-10)-L-(11) open reading frame. **(a)** Diagram of the sfGFPapa reporter and predicted transcription and processing steps resulting from treatment with the splicing inhibitor isoginkgetin. **(b)** COS7 cells transiently transfected with the sfGFPapa reporter for 24 hours were treated with either DMSO or isoginkgetin. Cells treated with isoginkgetin resulted in detectable green fluorescence after 24 hours. Scale bars in (**b**), 100 μm.

**Supplementary Figure 3.**
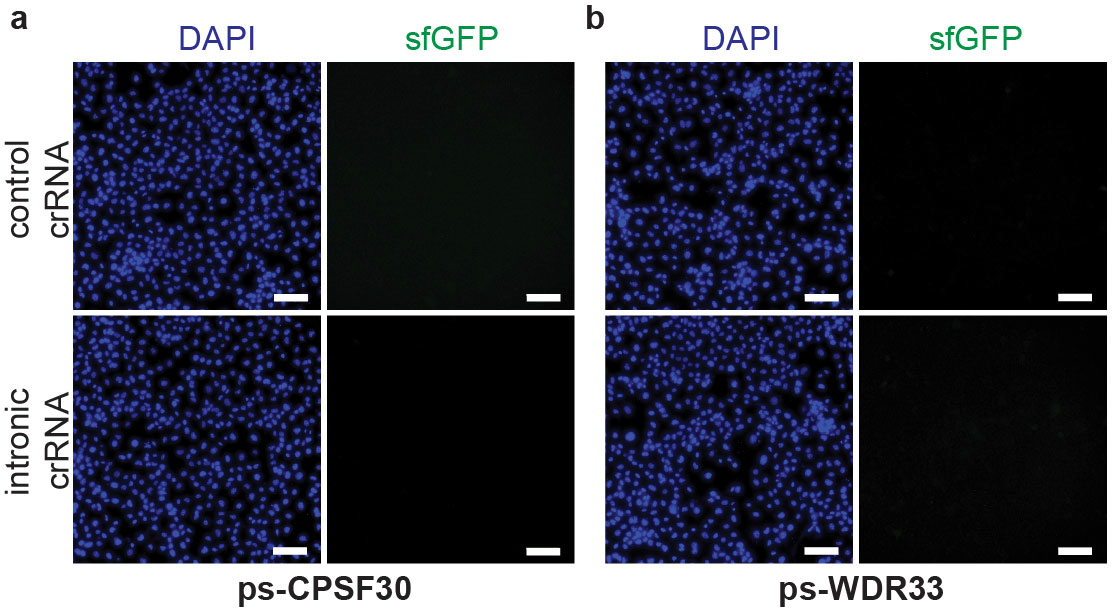
The dPspCas13b fusions to CPSP30 or WDR33 were not sufficient to promote cleavage and polyadenylation of the sfGFPapa reporter mRNA. (**a**) *Postscriptr* targeting of the sfGFPapa reporter using the dPspCas13b-CPSF30 or (**b**) – WDR33 fusion proteins using the sfGFPapa intronic-targeting crRNA did not result in detectable green fluorescence relative to a control non-targeting crRNA. Scale bars in (**a**) and (**b**), 100 μm.

**Supplementary Figure 4.**
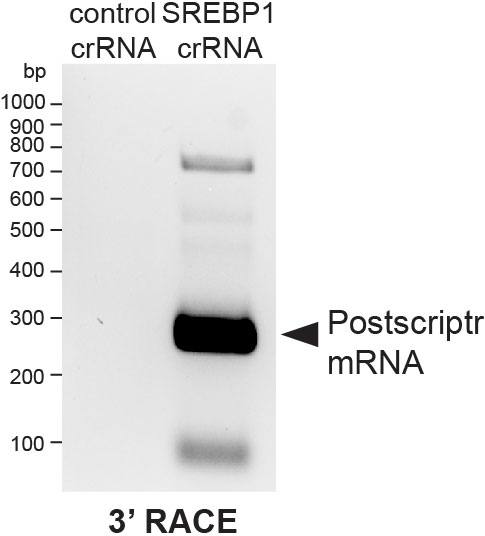
*Postscriptr*-mediated alternative cleavage and polyadenylation of human SREBP1 transcripts. PCR amplified 3’RACE products from cells expressing dPspCas13b-NUDT21 with non-targeting control or *SREBP1*-targeting crRNAs.

**SUPPLEMENTARY TABLE 1:**
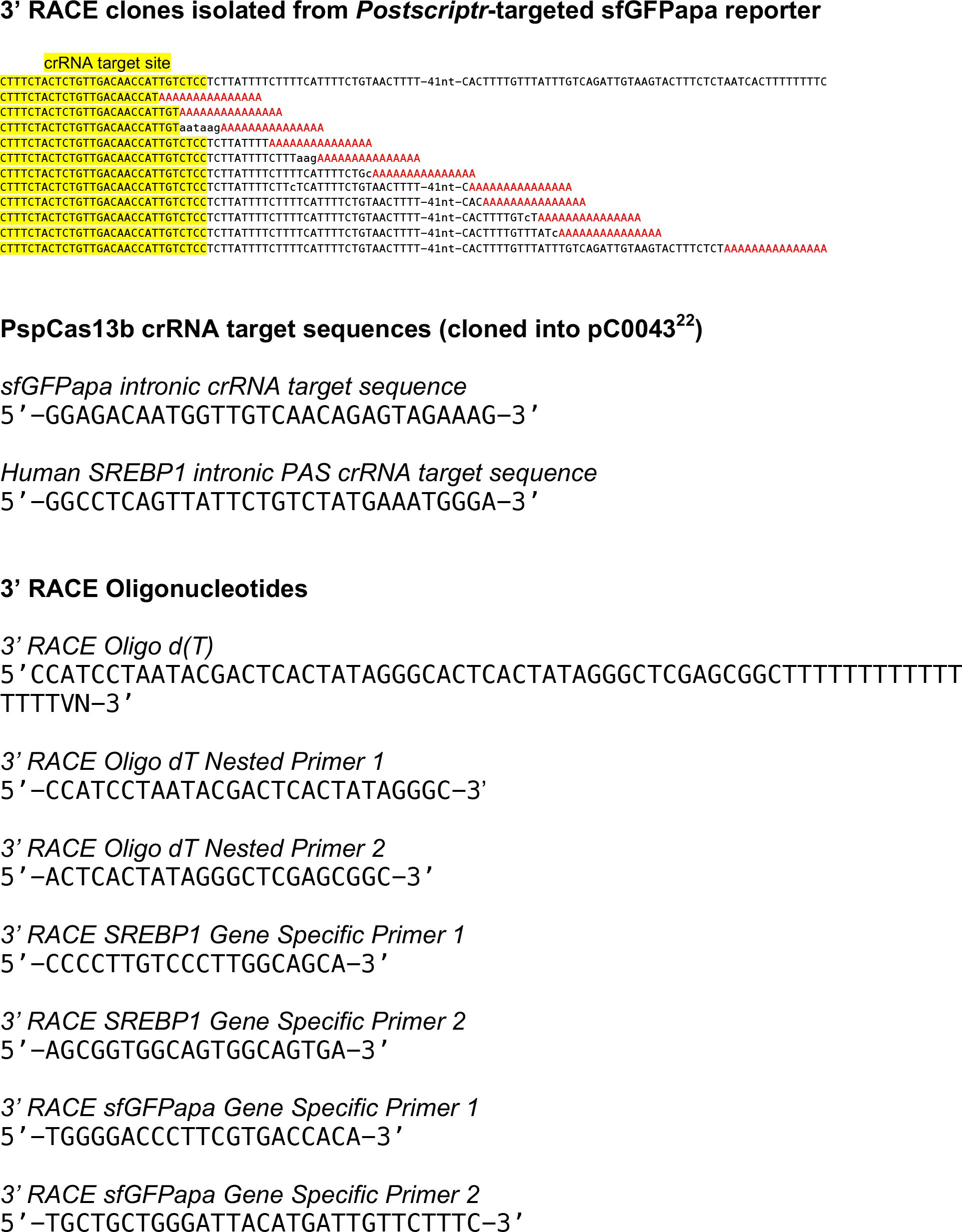
Sequences

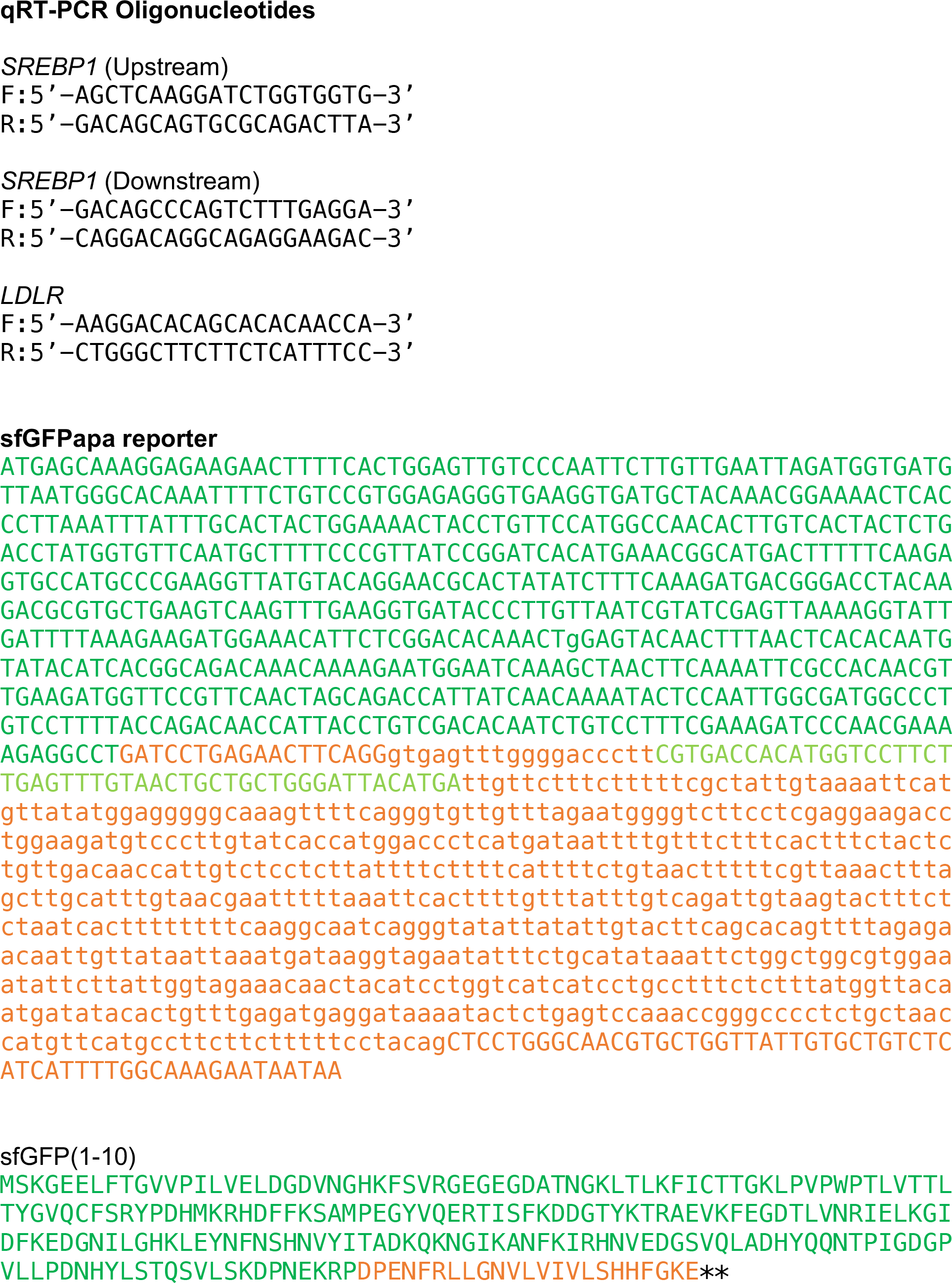

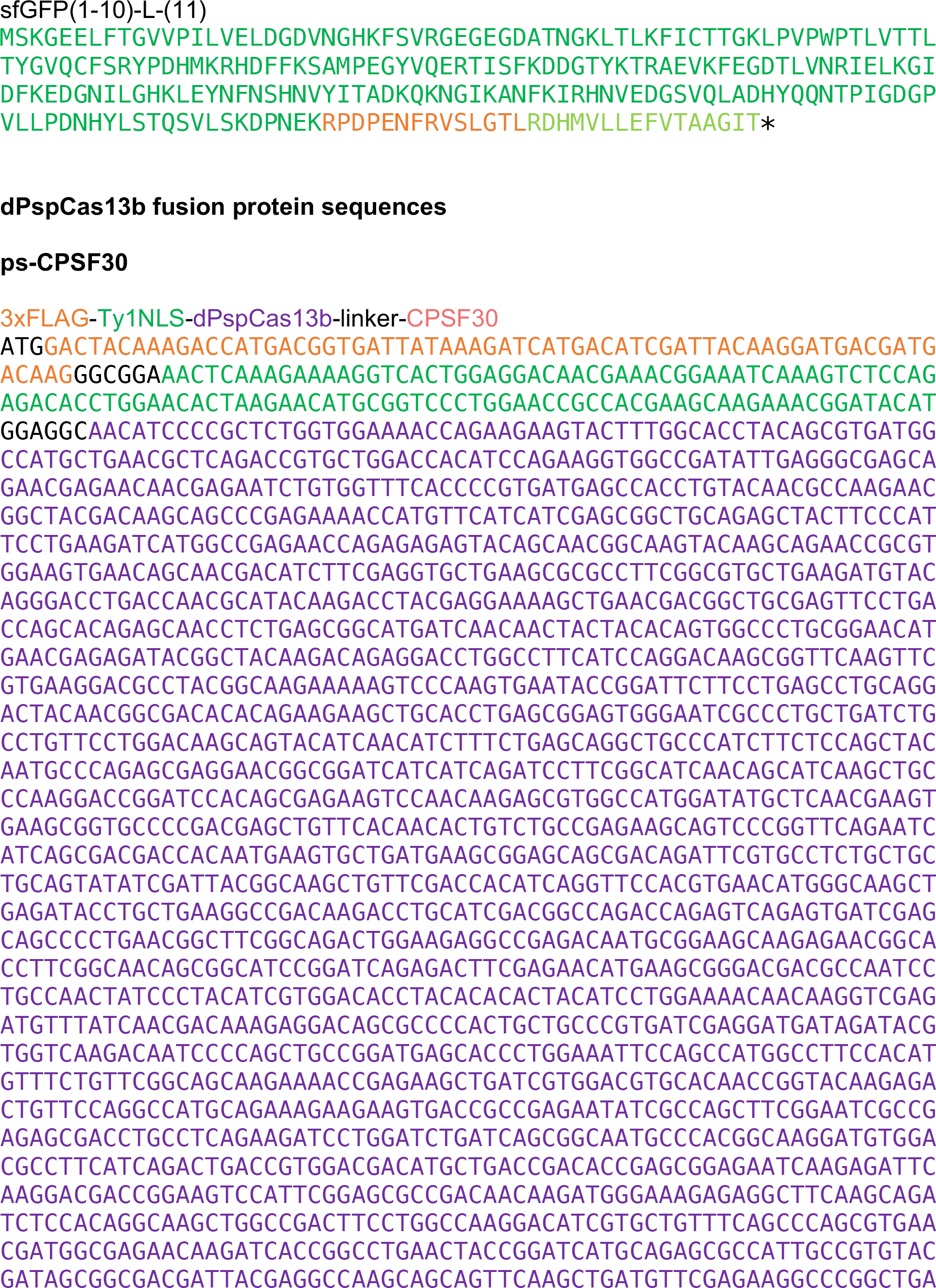

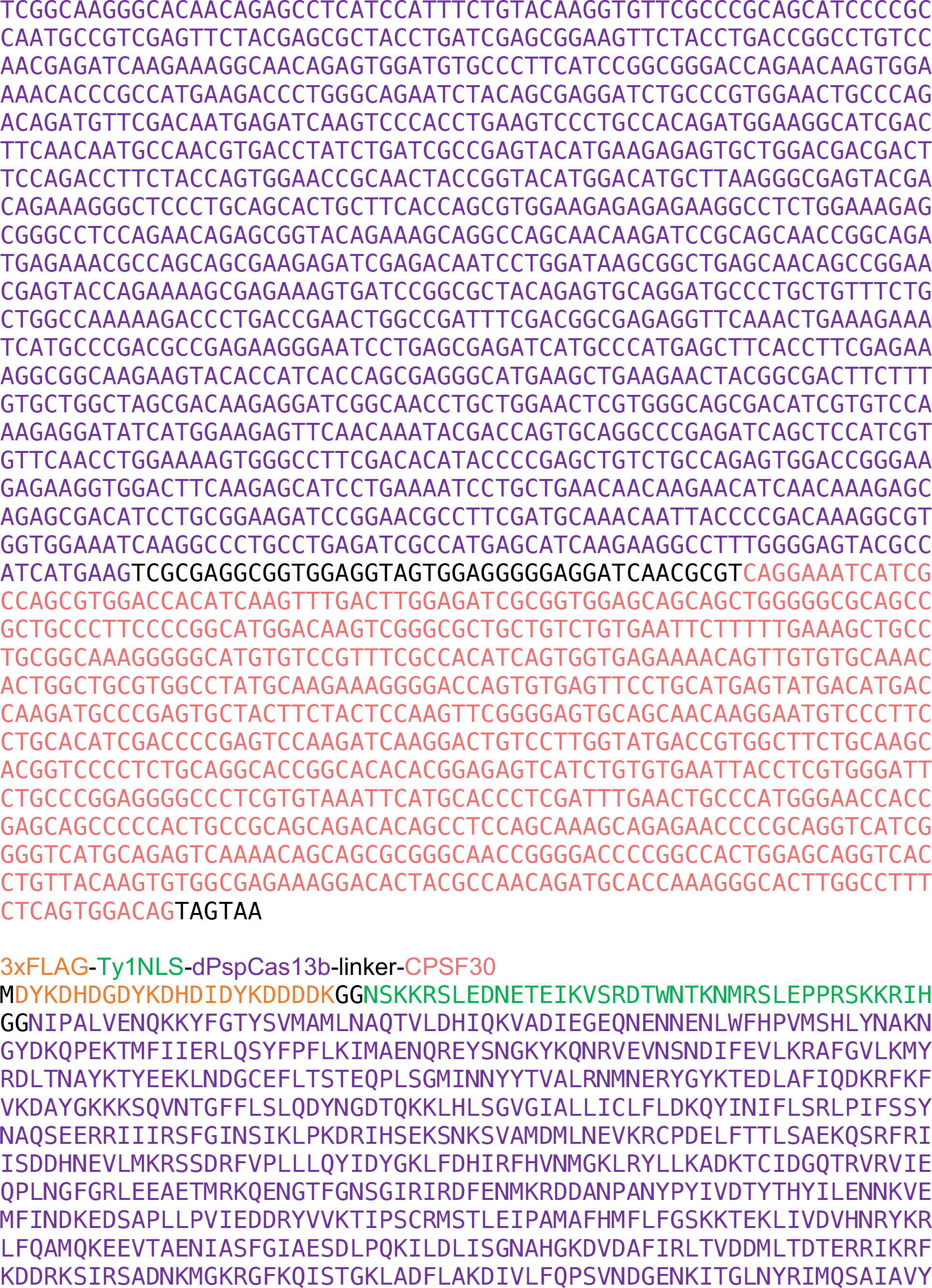

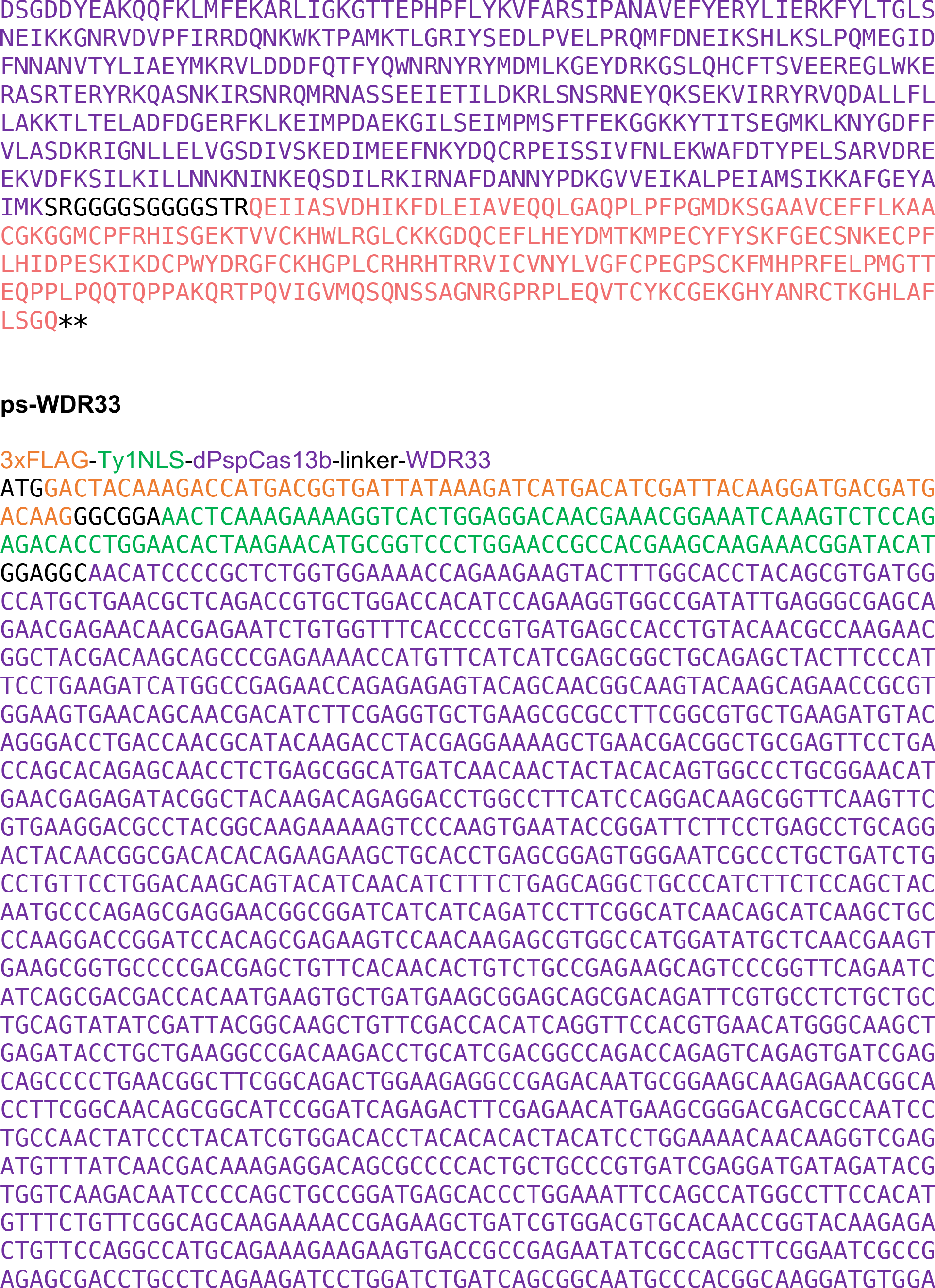

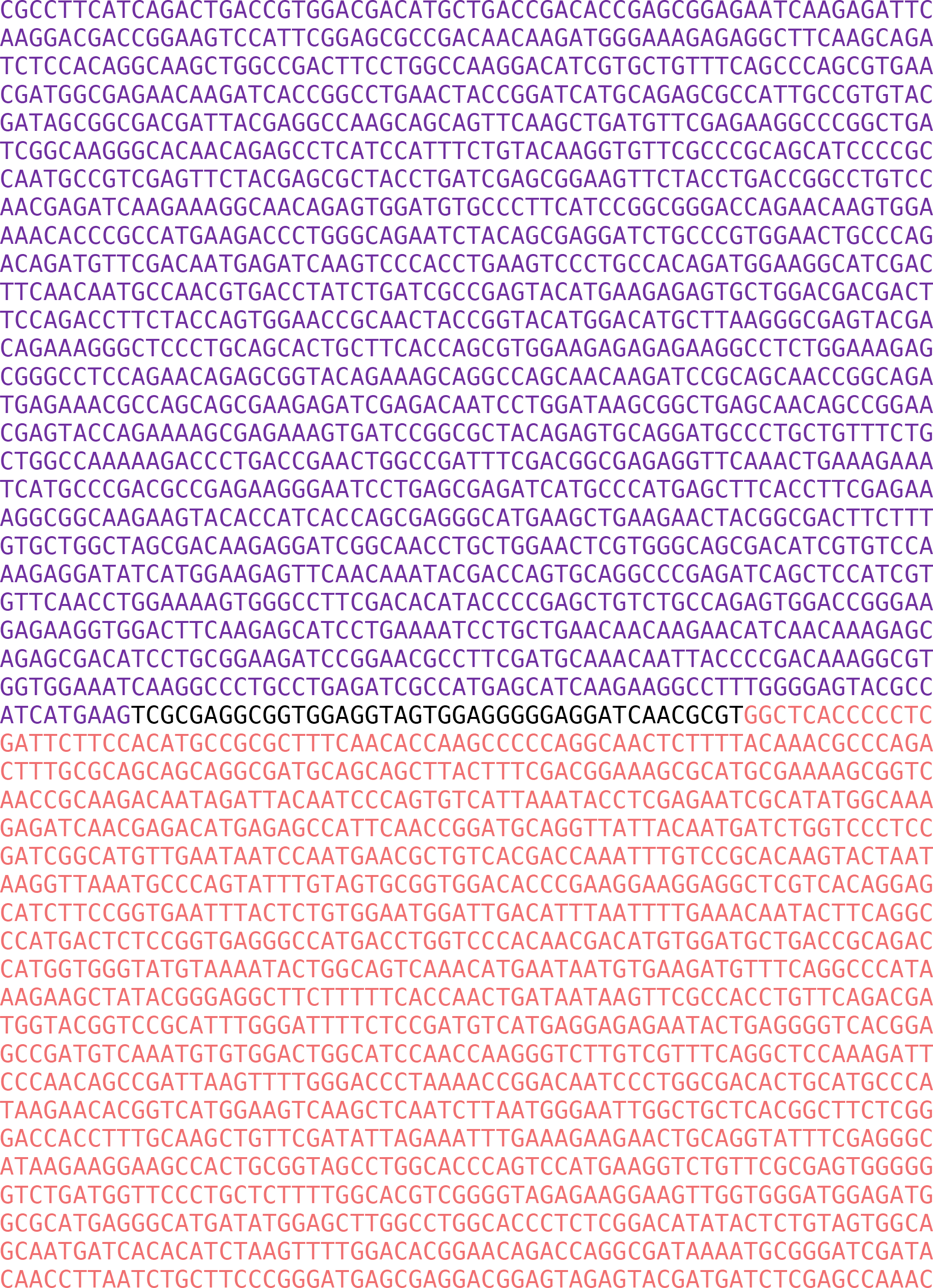

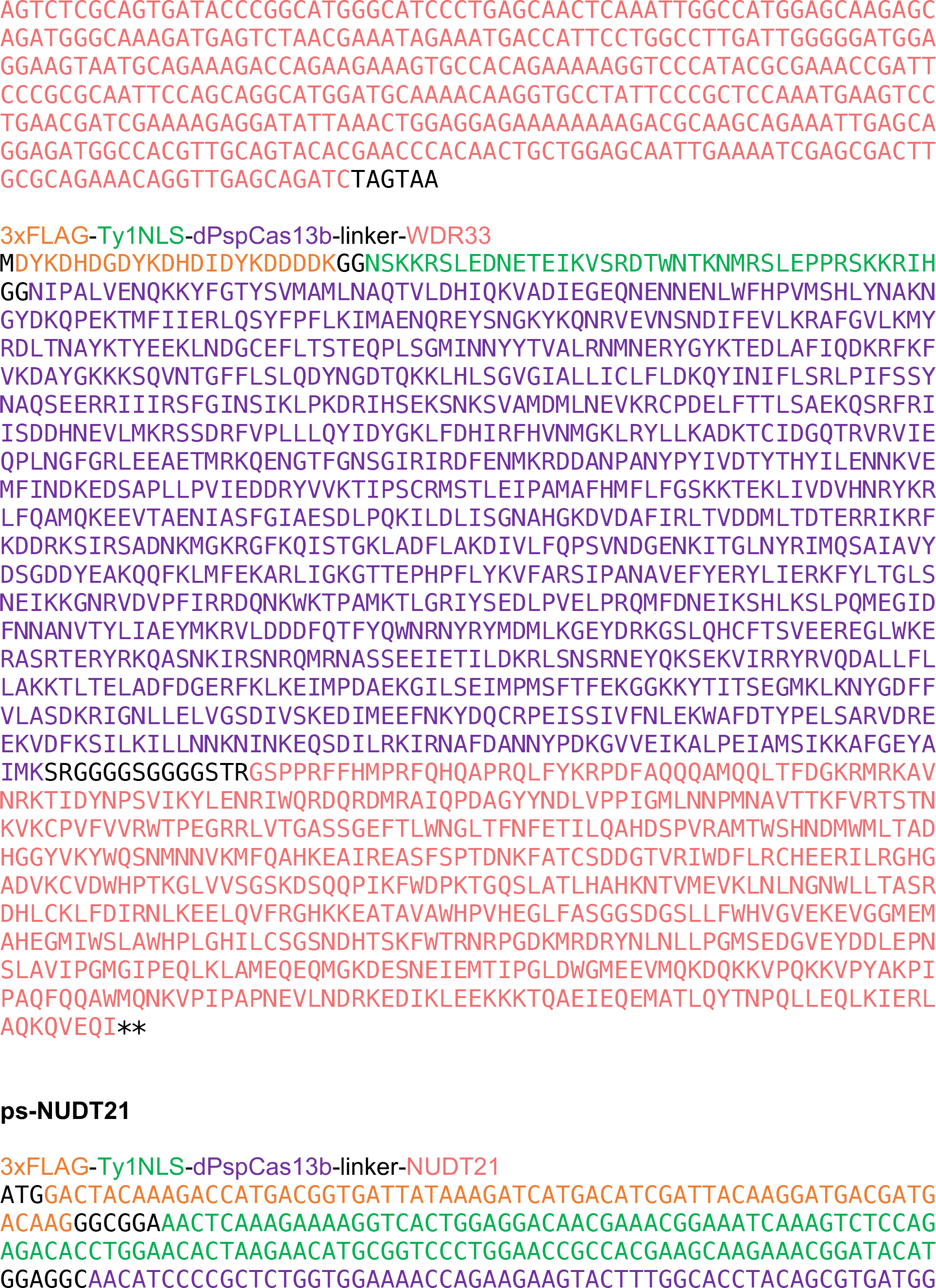

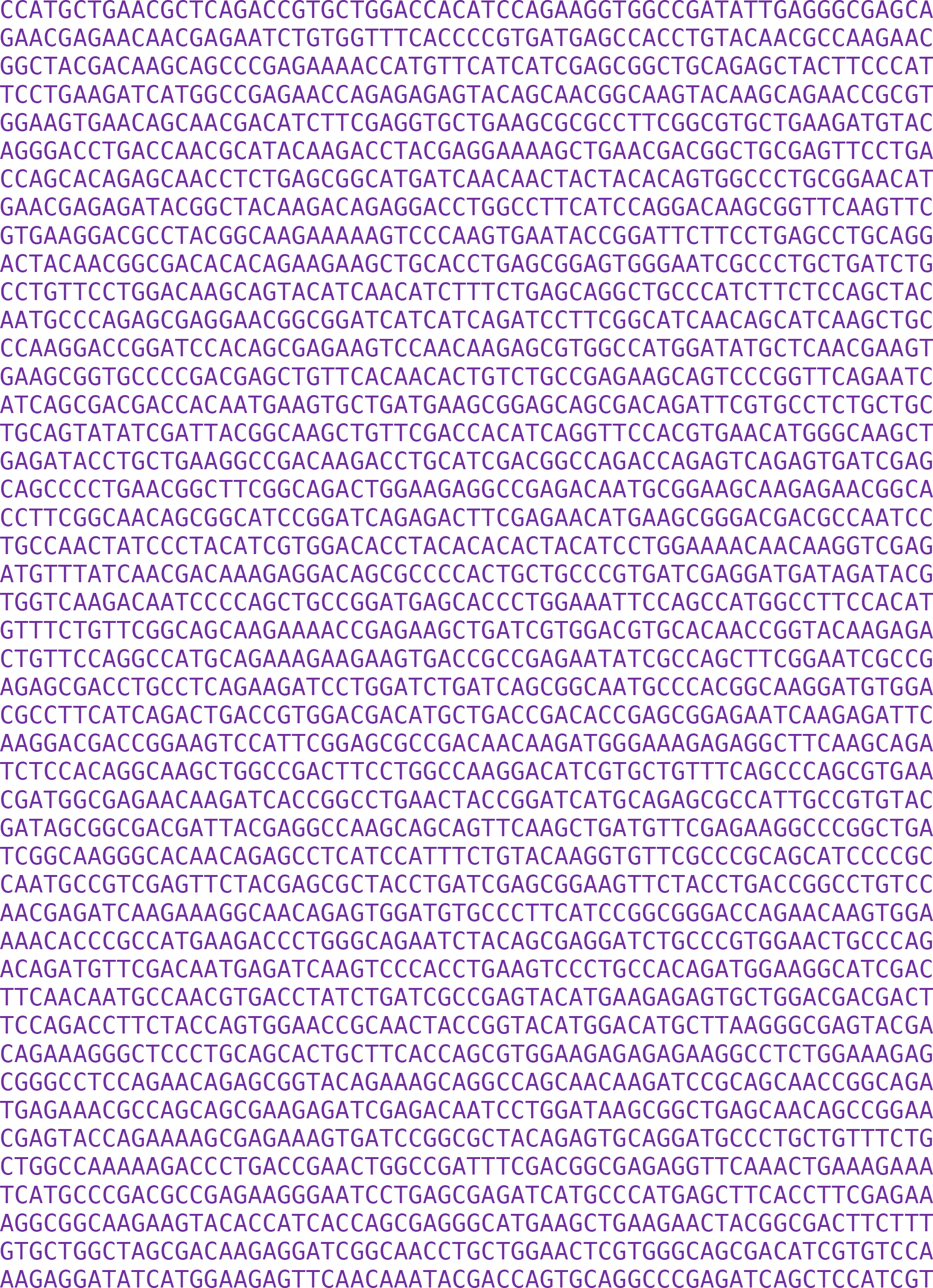

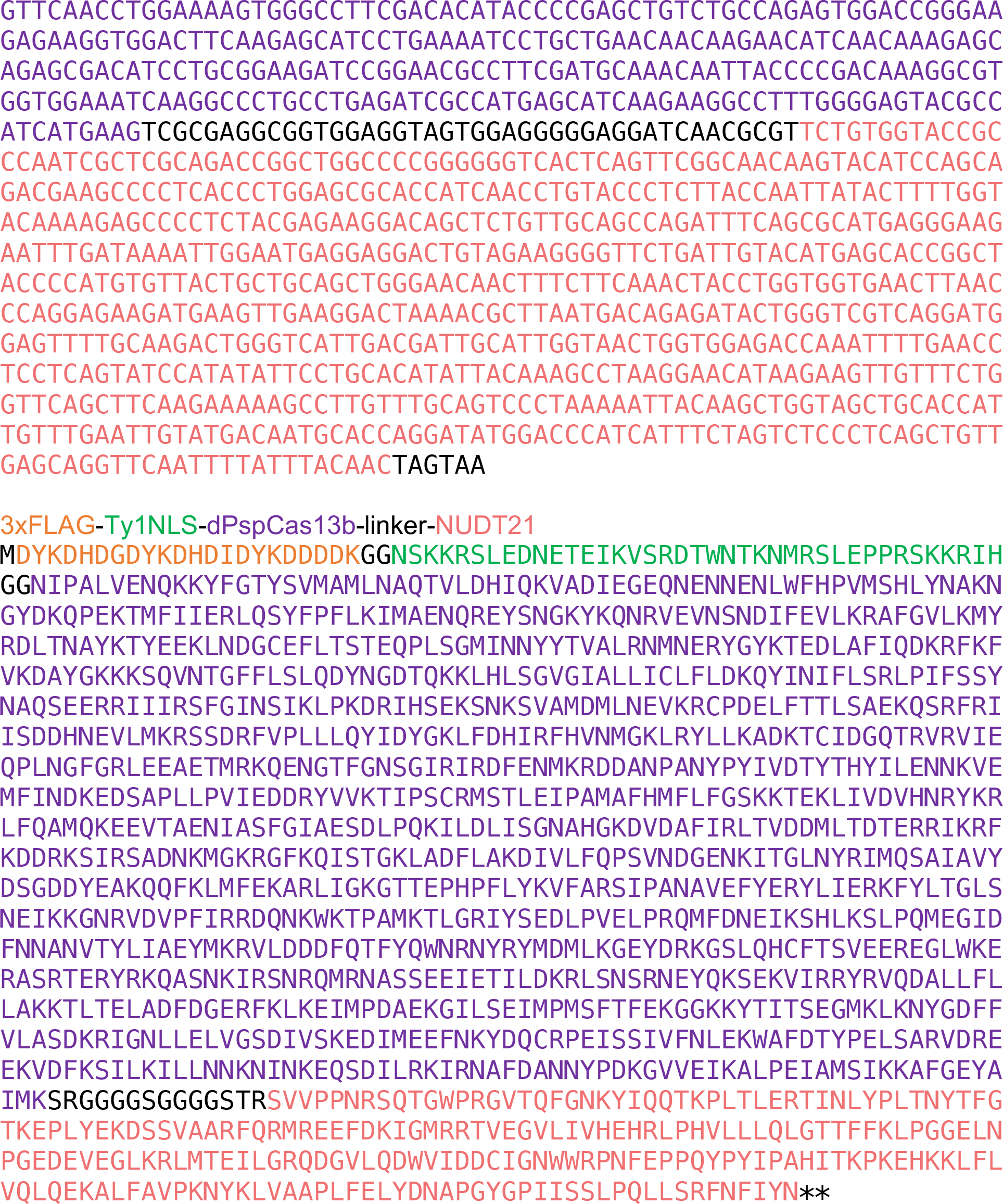

## REFERENCES

1 Mandel, C. R., Bai, Y. & Tong, L. Protein factors in pre-mRNA 3’-end processing. Cell Mol Life Sci 65, 1099–1122, doi:10.1007/s00018-007-7474-3 (2008).

2 Xiang, K., Tong, L. & Manley, J. L. Delineating the structural blueprint of the pre-mRNA 3’-end processing machinery. Mol Cell Biol 34, 1894–1910, doi:10.1128/MCB.00084-14 (2014).

3 Colgan, D. F. & Manley, J. L. Mechanism and regulation of mRNA polyadenylation. Genes Dev 11, 2755–2766 (1997).

4 Guhaniyogi, J. & Brewer, G. Regulation of mRNA stability in mammalian cells. Gene 265, 11–23 (2001).

5 Wu, X. & Brewer, G. The regulation of mRNA stability in mammalian cells: 2.0. Gene 500, 10–21, doi:10.1016/j.gene.2012.03.021 (2012).

6 Carmody, S. R. & Wente, S. R. mRNA nuclear export at a glance. J Cell Sci 122, 1933–1937, doi:10.1242/jcs.041236 (2009).

7 Tian, B. & Manley, J. L. Alternative polyadenylation of mRNA precursors. Nat Rev Mol Cell Biol 18, 18–30, doi:10.1038/nrm.2016.116 (2017).

8 Elkon, R., Ugalde, A. P. & Agami, R. Alternative cleavage and polyadenylation: extent, regulation and function. Nat Rev Genet 14, 496–506, doi:10.1038/nrg3482 (2013).

9 Chang, J. W., Yeh, H. S. & Yong, J. Alternative Polyadenylation in Human Diseases. Endocrinol Metab (Seoul) 32, 413–421, doi:10.3803/EnM.2017.32.4.413 (2017).

10 Curinha, A., Oliveira Braz, S., Pereira-Castro, I., Cruz, A. & Moreira, A. Implications of polyadenylation in health and disease. Nucleus 5, 508–519, doi:10.4161/nucl.36360 (2014).

11 Tian, B. & Graber, J. H. Signals for pre-mRNA cleavage and polyadenylation. Wiley Interdiscip Rev RNA 3, 385–396, doi:10.1002/wrna.116 (2012).

12 Chan, S. L. et al. CPSF30 and Wdr33 directly bind to AAUAAA in mammalian mRNA 3’ processing. Genes Dev 28, 2370–2380, doi:10.1101/gad.250993.114 (2014).

13 Venkataraman, K., Brown, K. M. & Gilmartin, G. M. Analysis of a noncanonical poly(A) site reveals a tripartite mechanism for vertebrate poly(A) site recognition. Genes Dev 19, 1315–1327, doi:10.1101/gad.1298605 (2005).

14 Zhu, Y. et al. Molecular Mechanisms for CFIm-Mediated Regulation of mRNA Alternative Polyadenylation. Mol Cell 69, 62–74 e64, doi:10.1016/j.molcel.2017.11.031 (2018).

15 Ruegsegger, U., Blank, D. & Keller, W. Human pre-mRNA cleavage factor Im is related to spliceosomal SR proteins and can be reconstituted in vitro from recombinant subunits. Mol Cell 1, 243–253 (1998).

16 Brown, K. M. & Gilmartin, G. M. A mechanism for the regulation of pre-mRNA 3’ processing by human cleavage factor Im. Mol Cell 12, 1467–1476 (2003).

17 Zhao, J., Hyman, L. & Moore, C. Formation of mRNA 3’ ends in eukaryotes: mechanism, regulation, and interrelationships with other steps in mRNA synthesis. Microbiol Mol Biol Rev 63, 405–445 (1999).

18 Abudayyeh, O. O. et al. C2c2 is a single-component programmable RNA-guided RNA-targeting CRISPR effector. Science 353, aaf5573, doi:10.1126/science.aaf5573 (2016).

19 Smargon, A. A. et al. Cas13b Is a Type VI-B CRISPR-Associated RNA-Guided RNase Differentially Regulated by Accessory Proteins Csx27 and Csx28. Mol Cell 65, 618–630 e617, doi:10.1016/j.molcel.2016.12.023 (2017).

20 East-Seletsky, A., O’Connell, M. R., Burstein, D., Knott, G. J. & Doudna, J. A. RNA Targeting by Functionally Orthogonal Type VI-A CRISPR-Cas Enzymes. Mol Cell 66, 373–383 e373, doi:10.1016/j.molcel.2017.04.008 (2017).

21 Abudayyeh, O. O. et al. RNA targeting with CRISPR-Cas13. Nature 550, 280–284, doi:10.1038/nature24049 (2017).

22 Cox, D. B. T. et al. RNA editing with CRISPR-Cas13. Science 358, 1019–1027, doi:10.1126/science.aaq0180 (2017).

23 O’Connell, M. R. Molecular Mechanisms of RNA Targeting by Cas13-containing Type VI CRISPR-Cas Systems. J Mol Biol 431, 66–87, doi:10.1016/j.jmb.2018.06.029 (2019).

24 McLane, L. M., Pulliam, K. F., Devine, S. E. & Corbett, A. H. The Ty1 integrase protein can exploit the classical nuclear protein import machinery for entry into the nucleus. Nucleic Acids Res 36, 4317–4326, doi:10.1093/nar/gkn383 (2008).

25 Kenna, M. A., Brachmann, C. B., Devine, S. E. & Boeke, J. D. Invading the yeast nucleus: a nuclear localization signal at the C terminus of Ty1 integrase is required for transposition in vivo. Mol Cell Biol 18, 1115–1124 (1998).

26 Kamiyama, D. et al. Versatile protein tagging in cells with split fluorescent protein. Nat Commun 7, 11046, doi:10.1038/ncomms11046 (2016).

27 Feng, S. et al. Improved split fluorescent proteins for endogenous protein labeling. Nat Commun 8, 370, doi:10.1038/s41467-017-00494-8 (2017).

28 Pedelacq, J. D., Cabantous, S., Tran, T., Terwilliger, T. C. & Waldo, G. S. Engineering and characterization of a superfolder green fluorescent protein. Nat Biotechnol 24, 79–88, doi:10.1038/nbt1172 (2006).

29 O’Brien, K., Matlin, A. J., Lowell, A. M. & Moore, M. J. The biflavonoid isoginkgetin is a general inhibitor of Pre-mRNA splicing. J Biol Chem 283, 33147–33154, doi:10.1074/jbc.M805556200 (2008).

30 Jin, Y. & Bian, T. Nontemplated nucleotide addition prior to polyadenylation: a comparison of Arabidopsis cDNA and genomic sequences. RNA 10, 1695–1697, doi:10.1261/rna.7610404 (2004).

31 Horton, J. D., Goldstein, J. L. & Brown, M. S. SREBPs: activators of the complete program of cholesterol and fatty acid synthesis in the liver. J Clin Invest 109, 1125–1131, doi:10.1172/JCI15593 (2002).

32 Brown, M. S. & Goldstein, J. L. The SREBP pathway: regulation of cholesterol metabolism by proteolysis of a membrane-bound transcription factor. Cell 89, 331–340 (1997).

33 Brown, M. S., Ye, J., Rawson, R. B. & Goldstein, J. L. Regulated intramembrane proteolysis: a control mechanism conserved from bacteria to humans. Cell 100, 391–398 (2000).

34 Kim, H. & Lee, Y. Interaction of poly(A) polymerase with the 25-kDa subunit of cleavage factor I. Biochem Biophys Res Commun 289, 513–518, doi:10.1006/bbrc.2001.5992 (2001).

35 Brumbaugh, J. et al. Nudt21 Controls Cell Fate by Connecting Alternative Polyadenylation to Chromatin Signaling. Cell 172, 106–120 e121, doi:10.1016/j.cell.2017.11.023 (2018).

36 Sartini, B. L., Wang, H., Wang, W., Millette, C. F. & Kilpatrick, D. L. Pre-messenger RNA cleavage factor I (CFIm): potential role in alternative polyadenylation during spermatogenesis. Biol Reprod 78, 472–482, doi:10.1095/biolreprod.107.064774 (2008).

37 Levitt, N., Briggs, D., Gil, A. & Proudfoot, N. J. Definition of an efficient synthetic poly(A) site. Genes Dev 3, 1019–1025 (1989).

38 Anderson, K. M. et al. Transcription of the non-coding RNA upperhand controls Hand2 expression and heart development. Nature 539, 433–436, doi:10.1038/nature20128 (2016).

39 Joung, J. et al. Genome-scale activation screen identifies a lncRNA locus regulating a gene neighbourhood. Nature 548, 343–346, doi:10.1038/nature23451 (2017).

40 Engreitz, J. M. et al. Local regulation of gene expression by lncRNA promoters, transcription and splicing. Nature 539, 452–455, doi:10.1038/nature20149 (2016).

41 Rinn, J. L. & Chang, H. Y. Genome regulation by long noncoding RNAs. Annu Rev Biochem 81, 145–166, doi:10.1146/annurev-biochem-051410-092902 (2012).

42 Higgs, D. R. et al. Alpha-thalassaemia caused by a polyadenylation signal mutation. Nature 306, 398–400 (1983).

43 Orkin, S. H., Cheng, T. C., Antonarakis, S. E. & Kazazian, H. H., Jr. Thalassemia due to a mutation in the cleavage-polyadenylation signal of the human beta-globin gene. EMBO J 4, 453–456 (1985).

44 Rund, D. et al. Two mutations in the beta-globin polyadenylylation signal reveal extended transcripts and new RNA polyadenylylation sites. Proc Natl Acad Sci U S A 89, 4324–4328 (1992).

